# Ligand-specific effects of 5-HT2A receptor antagonists on fear extinction in C57BL/6J mice: Comparative insights from MDL 11,939 and MDL 100,907

**DOI:** 10.64898/2026.06.29.735330

**Authors:** Anastasia Tyulmenkova, Robert W. Stackman

**Author notes:** To whom correspondence should be addressed: Robert W. Stackman Jr., Ph.D., Florida Atlantic University, John D. MacArthur Campus, Jupiter, FL 33458. USA.

## Abstract

Serotonin (5-HT) 2A receptors (5-HT_2_AR) modulate corticolimbic circuits regulating fear extinction. Although activation of these receptors has been shown to facilitate fear extinction, the behavioral consequences of 5-HT_2_AR antagonism during extinction is not well defined. Here, we examined the systemic effects of two 5-HT_2_A receptor antagonists, the mixed 5-HT_2_A/_2_C antagonist MDL 11,939 (Glemanserin) and the selective 5-HT_2_A antagonist MDL 100,907 (Volinanserin) on fear extinction in adult C57BL/6J mice. Prior to drug administration, mice assigned to future treatment groups acquired comparable conditioned freezing responses during delay fear conditioning. Twenty-four hours later, acute administration of MDL 11,939 (1.0 mg/kg) or MDL 100,907 (0.01 mg/kg) increased freezing to the first conditioned stimulus (CS) presentation on Extinction Day 1, indicating enhanced expression of conditioned fear. However, acquisition of fear extinction differed between the respective cohorts of mice treated with the two 5-HT_2_AR antagonists. Repeated administration of MDL 11,939 significantly impaired extinction, as evidenced by increased freezing across extinction trials and an increased number of trials required to reach extinction criterion. In contrast, MDL 100,907 has reported affinity for did not significantly alter extinction under either acute or repeated dosing conditions. Because MDL 11,939 has reported affinity for 5-HT_2_C receptors, we tested potential contributions of 5-HT_2_C receptor antagonism in a separate cohort of mice using two doses of the selective 5-HT_2_C antagonist, SB 242084. Neither dose affected conditioned fear expression, extinction learning, or trials required to reach extinction criterion. Together, these findings demonstrate ligand-specific and dose-dependent effects of 5-HT_2_AR antagonism on fear extinction and suggest that distinct intracellular receptor signaling pathways may differentially regulate extinction-related behavior.

**Highlights:**

- MDL 11,939 and MDL 100,907 differentially affect fear extinction behavior
- Repeated MDL 11,939 impairs extinction learning in C57BL/6J mice
- MDL 100,907 did not significantly alter extinction learning
- Selective 5-HT_2_C antagonism does not influence extinction
- Ligand-dependent 5-HT2A signaling shapes extinction outcomes

## Introduction

Pathological fear has accompanied human experiences across recorded history, with descriptions of trauma related symptoms appearing in ancient texts long before modern psychiatry (Abdul-Hamid and Hughes, 2014). These historical recordings underscore the biological challenge of suppressing fear memories once an adverse experience has been established in memory. However, when fear becomes persistent or is triggered by innocuous cues, it can manifest as maladaptive anxiety, avoidance, or post-traumatic stress. Through associative learning, neutral environmental cues can acquire emotional salience by being paired with an aversive event, such that these cues later evoke defensive behaviors even in the absence of danger. Laboratory animal and human studies have utilized Pavlovian contextual- and cue-based fear conditioning protocols to investigate the mechanisms of fear memory processes. These protocols allow precise measurement of how fear memories are encoded, consolidated, expressed and suppressed. Once a conditioned fear memory is acquired, repeated exposure to a threat-associated context or cue in the absence of aversive stimuli can lead to a gradual reduction in the fear response strength over time, a phenomenon referred to as “fear extinction” (Myers et al., 2006). Fear extinction is defined as the reduction in a conditioned fear response (CR) that occurs following repeated, nonreinforced presentations of a conditioning stimulus (CS) that was previously paired with an aversive unconditioned stimulus (US). Research spanning from as far back as 1970 has theorized that extinction is not a process of a memory being erased, but instead the acquisition of an inhibitory memory, which acts to suppress the strength of the fear CR (Bouton et al., 2021). The initial memory of the CS-US fear association is intact, yet the new inhibitory memory extinguishes the CR elicited by the CS. Recently, another theory proposes that not only does the inhibitory memory need to be intact, but extinction can be further driven by activated dopaminergic projections from the ventral tegmental area to the basolateral amygdala (BLA) reward pathway to enhance extinction learning (Zhang et al., 2025).

### Neural circuits of fear extinction

Fear conditioning and extinction are mediated by a distributed neural network centered on the amygdala, medial prefrontal cortex (mPFC), and hippocampus. At the core of the fear conditioning network is the basolateral amygdala (BLA), which serves as a critical hub for sensory integration and synaptic plasticity (Maren and Holmes, 2016). During fear conditioning, BLA neurons undergo synaptic plasticity triggered by the reception of converging excitatory inputs: CS-related signals via the auditory thalamus and cortex, and US-related signals from midbrain regions such as the parabrachial nucleus (Bouton et al., 2021). Upon integration of these signals, the BLA activates the central amygdala (CeA), which orchestrates the expression of defensive behaviors in response to the threatening stimulus. The BLA also projects to the mPFC, a region instrumental in modulating fear learning and extinction. Within the mPFC, the prelimbic (PL) cortex enhances fear expression, whereas the infralimbic (IL) cortex facilitates fear inhibition and the consolidation of fear extinction memory (Hill and Martinowich, 2016). Top-down inhibition of fear expression is exerted through IL projections to the BLA, which recruit intercalated GABAergic cells to suppress CeA output and, in turn, suppress fear expression. This extinction-associated plasticity in the IL is regulated by dopaminergic input from the ventral tegmental area, which is responsive to negative prediction errors. Specifically, instances when expected aversive outcomes fail to occur. Increased dopaminergic signaling in such cases activates D2 receptors in the IL, promoting extinction learning (Salinas-Hernández et al., 2018). Additionally, endogenous mu-opioid receptor signaling in the periaqueductal gray (PAG) supports extinction via modulation of IL neuronal excitability. Contextual regulation of fear extinction is facilitated by hippocampal inputs to the mPFC, particularly from the CA1 region. Hippocampal inputs enable discrimination between safe and threatening environments, thereby guiding appropriate fear responses (Bouton et al., 2021). Together, the amygdala, mPFC and hippocampus comprise a dynamic circuit that governs the encoding, consolidation, expression, and extinction of conditioned fear.

Despite the well-established roles of mPFC subregions in fear regulation, experimental outcomes following mPFC stimulation or inhibition via optogenetic, chemogenetic, and pharmacologic approaches have yielded inconsistent behavioral results. These discrepancies are likely due to methodological differences and the inherent complexity of threat processing within the mPFC. The PL and IL are anatomically proximate but functionally distinct; thus, slight variation in experimental parameters can differentially engage circuits supporting either fear expression or extinction (Alexandra Kredlow et al., 2022). At the molecular level, fear conditioning and extinction are associated with distinct gene expression patterns in the PL and IL, respectively. These changes occur through transcriptional and epigenetic mechanisms, prominently involving brain-derived neurotrophic factor (BDNF) expression, which is essential for long-term memory consolidation and synaptic remodeling. Importantly, these molecular processes are influenced by neuromodulatory systems, particularly serotonergic input from the dorsal raphe nucleus (DRN). The DRN projects to both the mPFC and BLA, where serotonin modulates neuronal excitability, synaptic plasticity, and gene expression (Sengupta and Holmes, 2019). Among the various serotonin receptors, the 5-HT_2_A receptor (5-HT_2_AR) has emerged as a key regulator of fear learning, memory, and extinction with a unique mechanism of action (Zhang and Stackman, 2015). Pharmacological activation of 5-HT_2_ARs facilitated both consolidation and extinction of fear memories in male C57BL/6J mice, suggesting a potential role for this receptor in modulating adaptive fear learning (Zhang et al., 2013). Facilitation of fear extinction by 5-HT_2_AR agonism may be linked to intracellular cascades involving phospholipase C (PLC) and calcium signaling, implicating pathways in the expression of BDNF and synaptic plasticity. These molecular events may serve to fine-tune activity within the mPFC-amygdalar circuits, potentially supporting extinction learning, though precise mechanisms remain unknown.

### Role of 5-HT_2_A receptors in fear extinction

Taken together, converging evidence supports the hypothesis that the 5-HT_2_AR plays a central role in regulating fear extinction, due to its dense expression in the mPFC, hippocampus and amygdala. Activation of 5-HT_2_ARs modulates neuronal excitability and initiates intracellular signaling cascades, primarily PLC and calcium-dependent pathways, which in turn enhance expression of plasticity-associated genes such as *BDNF* (Cummins et al., 2025; Raote et al., 2007; Vaidya et al., 1997). These molecular events facilitate synaptic remodeling required for extinction learning. Genetic models further support the role: deletion of 5-HT_2_ARs in 129S6/SvEv mice alters expression of immediate early and stress-responsive genes in the mPFC and hippocampus (Jaggar et al., 2017). These findings emphasize the 5-HT_2_AR’s contribution to stress-related learning and memory.

#### Functional Selectivity and Agonist Effects

The behavioral effects of 5-HT_2_AR activation are shaped by functional selectivity or biased agonism, whereby different ligands stabilize distinct receptor conformations, and engage divergent signaling cascades (López-Giménez and González-Maeso, 2018). Hallucinogenic agonists such as 2,5-dimethoxy-4-iodoamphetamine (DOI), psilocybin, and TCB-2 reliably facilitate fear extinction (Du et al., 2023; Pędzich et al., 2022; Woodburn et al., 2024; Zhang et al., 2013). For example, DOI increased c-Fos expression in the amygdala, enhanced prefrontal cortical high-frequency gamma oscillations linked to neural synchrony, and enhanced memory encoding, and consolidation (Carr et al., 2011; Gener et al., 2019; Pędzich et al., 2022). These mechanisms are thought to support the updating of maladaptive fear memories, thereby improving emotional regulation.

Recent evidence suggests that psychedelics may exert their unique effects by engaging intracellular 5-HT_2_AR pools that are less accessible to endogenous serotonin. Such compartmentalized receptor activation yields distinct intracellular signaling outcomes (Vargas et al., 2023). While non-hallucinogenic agonists (e.g., lisuride and ergotamine) primarily activate canonical PLC signaling, hallucinogenic agonists preferentially recruit β-arrestin-2-dependent pathways. Notably, several DOI-mediated responses persist in β-arrestin-2 knockout models, indicating the complexity of receptor-effector coupling (Schmid et al., 2008).

#### Antagonist Effects and Pharmacological Challenges

Although most research has emphasized agonism, 5-HT_2_AR antagonism also modulates extinction learning, with effects that are dose- and time-dependent. Selective antagonists, such as MDL 11,939 (also known as Glemanserin, GLE) and MDL 100,907 (Volinanserin, VOL) can impair fear extinction in some conditions but facilitate long-term consolidation in others (Dougherty and Aloyo, 2011; Hagsäter et al., 2021). In their review, Zhang and Stackman (2015) highlight that 5-HT_2_AR antagonism can reduce cognitive flexibility and disrupt fear-related learning processes, underscoring the receptor’s bidirectional influence on memory systems.

Early pharmacological studies demonstrated that serotonin acting at cortical 5-HT_2_ receptors stimulates phosphoinositide hydrolysis, and the potency of antagonists on this intracellular signal corresponds with their binding affinities (Conn and Sanders-Bush, 1985). These findings promoted subsequent investigations to distinguish the specific contributions of 5-HT_2_A, 5-HT_2_B and 5-HT_2_C receptor subtypes using putative selective antagonists (Casey et al., 2022). However, the interpretation of behavioral studies is complicated by the limited availability of 5-HT_2_ receptor ligands with absolute selectivity. For instance, MDL 11,939 binds both 5-HT_2_A and 5-HT_2_C receptors, confounding attribution of effects to a single receptor subtype.

### Selective 5-HT_2_A antagonists MDL 11,939 and MDL 100,907

5-HT_2_AR are constitutively active in neurons. Selective antagonists such as MDL 11,939 and MDL 100,907 can silence these receptors and potentially interfere with the acquisition of fear extinction (Johnson et al., 1996; Sullivan et al., 2015). In rabbits, repeated administration of MDL 11,939 and MDL 100,907 increased 5-HT_2_AR expression and accelerated the rate of learning response. Psychedelic agonists with both 5-HT_2_AR and 5-HT_2_CR activity decreased 5-HT_2_AR expression with no effect on 5-HT_2_CR expression. In contrast, both selective and nonselective 5-HT_2_A antagonists increased 5-HT_2_AR expression while decreasing 5-HT_2_CR expression (Aloyo et al., 2001).

Administration of D-2-bromolysergic acid diethylamide hydrogen tartrate (BOL), which binds to both 5-HT_2_AR and 5-HT_2_CR with similar affinity, reduced 5-HT_2_AR expression. Whereas MDL 11,939 selectively increased 5-HT_2_AR expression without altering 5-HT_2_CR expression (Aloyo et al., 2001). Studies in C57BL/6N mice revealed a low baseline 5-HT_2_CR expression and stable 5-HT_2_AR expression after repeated exposure of MDL 11,939. In contrast, the psychedelic compound DOI led to down-regulation of 5-HT_2_AR expression (Dougherty and Aloyo, 2011).

In humans, the pharmacological effects of a single dose of MDL 11,939 dissipated within several hours of administration (Dudley et al., 1988). After a single dose of MDL 100,907, brain receptor binding in humans remained stable at 24 hours following the 20 mg dose, but decreased by approximately 20% after the 10 mg dose; the plasma elimination half-life was estimated at 6.56 ± 1.56 hours (Gründer et al., 1997). Unlike genetic deletion of 5-HT_2_ARs, pharmacological silencing is transient and may induce compensatory upregulation of receptor expression. In addition, coadministration of selective antagonist MDL 100,907 or the inverse agonist pimavanserin with the selective serotonin reuptake inhibitor (SSRI), citalopram facilitated fear extinction in Sprague-Dawley rats, whereas administration of either substance alone was ineffective (Hagsäter et al., 2021). However, although the authors reported statistically significant effects, they were unable to consistently reproduce the facilitation of fear extinction produced by MDL 100,907 at the 0.03 mg/kg dose. MDL 100,907 also enhanced SSRI-induced antidepressant-like behavior in Sprague-Dawley rats when combined with fluoxetine (Marek et al., 2005). Taken together, these findings suggest that selective 5-HT_2_AR antagonists may modulate fear extinction without producing unwanted psychedelic effects.

#### Comparison with 5-HT_2_C Receptors

In contrast to 5-HT_2_ARs, antagonism of 5-HT_2_CRs appears to more consistently facilitate fear extinction and reduce anxiety-like behavior. Selective 5-HT_2_CR antagonists, such as RS-102221 and SB 242084 (Bonhaus et al., 1997; Kennett et al., 1997) promote extinction, potentially by disinhibiting stress-regulatory circuits within the extended amygdala and prefrontal cortex (Minshall et al., 2024; Süß et al., 2022). These findings highlight divergent roles for these receptor subtypes, with 5-HT_2_ARs exerting bidirectional context-dependent influences on extinction, whereas 5-HT_2_CRs appear to serve a more unidirectional neuromodulatory role.

### Summary

Collectively, the available evidence indicates that 5-HT_2_AR signaling exerts complex, bidirectional effects on fear extinction. Agonism generally enhances extinction, likely through effects on synaptic plasticity and circuit synchrony, whereas antagonism produces more heterogeneous and context-dependent outcomes. In contrast, 5-HT_2_CR antagonism appears to reliably facilitate extinction, underscoring the importance of carefully delineating receptor subtype-specific contributions to fear regulation. Clarifying these mechanisms will be essential for translating preclinical findings into targeted interventions for fear-related psychopathologies. In the present study, we examined the effects of 5-HT_2_AR antagonists on the extinction of conditioned fear memory, building on our previous work on the modulatory role of 5-HT_2_AR in learning and memory processes in mice.

## 2. Methods

Fear conditioning and extinction procedures were adapted from (Zhang et al., 2013), using the same apparatus stimulus parameters. However, in the present study, we introduced pharmacological treatments with MDL 11,939 and MDL 100,907 prior to extinction training to assess their effects on fear memory modulation.

### 2.1 Animals

Adult male C57BL/6J mice (8–12 weeks old) were obtained from Jackson Laboratories (Bar Harbor, ME). Mice were group-housed (4 per cage) in polycarbonate cages with *ad libitum* access to food and water. The colony room was maintained under controlled temperature and humidity conditions with a 12-hour light/dark cycle (lights on at 7:30 AM). Bedding was changed before behavioral training commenced or delayed until after experimental completion. All behavioral testing was conducted during the light phase. All procedures adhered to the *National Institutes of Health Guide for the Care and Use of Laboratory Animals* and received approval by the Institutional Animal Care and Use Committee at Florida Atlantic University.

### 2.2 Apparatus

#### 2.2.1 Fear Conditioning and Extinction System for MDL 11,939 and MDL 100,907 Cohorts

Mice underwent fear conditioning and extinction using the MED Associates Near-Infrared Video Fear Conditioning System (Georgia, VT), following previously established protocols (Zhang et al., 2013). The system included four identical conditioning chambers (30.5 cm × 24.1 cm × 21 cm) featuring stainless steel grid floors that delivered scrambled foot shocks. Freezing behavior was automatically analyzed using the MED Associates software, which classified freezing as a period of immobility lasting more than 0.6 seconds, with fewer than 20 pixels of movement detected across consecutive video frames. Data were captured at 30 frames per second (fps), and freezing episodes were computationally identified when motion remained below the 20-pixel threshold for at least 18 frames (Zhang et al., 2013).

### 2.3 Habituation

For one week, mice were habituated daily to the experimental room and polycarbonate holding cages along with handling and i.p. injections, to acclimate them to drug-administration procedures.

### 2.4 Fear Conditioning and Extinction

#### 2.4.1 Delay Fear Conditioning and Context Exposure

Fear conditioning was modified from our previously reported protocol (Zhang et al., 2013), by adding pre-exposure to both Context A and Context B before conditioning. On the pre-exposure day, mice were allowed 5 min of free exploration in Context A and then were immediately placed in Context B for an additional 5 min. This procedure assessed baseline exploratory behavior and facilitated context discrimination prior to conditioning. On the following day, mice underwent delay fear conditioning in Context A. After a 60-s baseline period, mice received three CS-US pairings consisting of a 30-s tone (CS; 90 dB, 5 kHz) that co-terminated with a 1-s, 0.5-mA foot shock (US). Mice remained in Context A for 60 s following the final CS–US pairing before being returned to their home cages.

#### 2.4.2 Fear Extinction Training and Testing

To more thoroughly assess extinction learning, the protocol was modified from our previously reported single-session design (Zhang et al., 2013) to include three daily extinction sessions. On Extinction Day 1, mice were placed in Context B, which differed from Context A in lighting, texture, color, and odor. After a 60-s acclimation period, mice received 20 non-reinforced presentations of the CS, each separated by a 5-s intertrial interval (ITI). Identical extinction sessions were conducted in Context B on the following two days (Extinction Days 2 and 3). To control for generalized freezing unrelated to CS presentation, mice exhibiting pre-CS freezing levels >20% on Extinction Day 1 were excluded from subsequent analyses.

### 2.5 Pharmacological Agents and Experimental Design

To investigate the effects of 5-HT_2_AR antagonism on fear extinction, two selective antagonists were tested: MDL 11,939 (Glemanserin, GLE); *N-[4-(4-fluorobenzyl)-2-(4-methyl-piperazin-1-yl) -phenyl]-2-methylpropanamide*, Tocris Bioscience) and MDL 100,907 (Volinanserin, VOL) 7; *R-(+)-α-(2,3-dimethoxyphenyl)-1-[2-(4-fluorophenyl)ethyl]-4-piperidine-methanol*, Sigma-Aldrich). MDL 11,939 has been shown to influence fear-related learning and memory (Zhang et al., 2013), whereas MDL 100,907 has been widely used to examine serotonergic contributions to extinction learning and cognitive flexibility (Boulougouris et al., 2008; Pędzich et al., 2022; Zhang et al., 2013).

Mice were randomly assigned to one of two administration protocols:

1. **Acute administration:** A single IP injection of MDL 11,939 (0.5 mg/kg or 1 mg/kg) or MDL 100,907 (0.01 mg/kg or 0.03 mg/kg) given 30 min prior to Extinction Day 1 to assess immediate effects on extinction learning.
2. **Repeated administration:** The same drug regimens were administered 30 min prior to Extinction Day 1 and prior to Extinction Day 2 to examine the impact of repeated exposure on extinction processes.

Control mice received vehicle injections (VEH, 5% DMSO in saline) on the same schedule as their drug-treated counterparts.

### 2.6 Behavioral Analysis

Freezing behavior was quantified across all phases of the experiment. For the contextual fear test session, freezing levels were measured during the full 5-min sessions in Context A and Context B and expressed as total % freezing. During conditioning, freezing was recorded during each of the three CS presentations and the subsequent 30-s post-tone intervals. For cued extinction, freezing responses were analyzed across each of the 20 CS presentations within each extinction session to track acquisition of extinction over time.

### 2.7 Statistical Analysis

A one-way ANOVA, with Dunnett’s post hoc test, was used to assess treatment group differences in freezing response to the first CS presentation on Extinction Day 1 as a measure of the strength of the conditioned fear memory. Freezing behavior during extinction training was analyzed using two-way repeated-measures (RM) ANOVA with CS presentation (within-subjects) and treatment group (between-subjects) as factors. Assumptions of normality and homogeneity of variance were verified prior to analysis. When sphericity was violated, the Geisser–Greenhouse correction was applied, and corrected degrees of freedom were reported. Significant main effects or interactions were followed by Tukey’s multiple comparisons test to identify pairwise group differences.

To further assess the acquisition of fear extinction, a measure of trials-to-extinction was determined for each mouse; defined as the number of consecutive CS presentations required for the mouse to reach extinction criterion, operationalized as four or more consecutive CSs with freezing reduced to ≤50% of the freezing level observed during the third CS-US pairing on conditioning day. Mice that did not reach criterion were assigned the maximum value of 60 trials. Trials-to-extinction were analyzed using a one-way ANOVA, with Dunnett’s post hoc test used to compare each treatment group against the vehicle.

The proportion of mice meeting extinction criterion across groups and trial days were compared using Fisher’s exact test. Statistical significance for all analyses was set at *p* < 0.05. All analyses were performed in GraphPad Prism (GraphPad Software, San Diego, CA).

## 3. Results

### 3.1 Fear Conditioning Across Groups

To confirm baseline equivalence across future treatment groups (VEH, *n*=23; GLE 0.5 mg/kg, *n*=22; GLE 1.0 mg/kg, *n*=20; VOL 0.01 mg/kg, *n*=25; VOL 0.03 mg/kg, *n*=25) freezing measures from the first 60 s of the conditioning session and each CS-US pairing interval (30s CS plus the 30s that followed). Fear conditioning data were analyzed separately for mice assigned to future MDL 11,939 and MDL 100,907 treatment groups.

In the MDL 11,939 cohort (VEH, GLE 0.5 mg/kg, GLE 1.0 mg/kg), a two-way RM ANOVA revealed a significant main effect of CS-US presentation on freezing behavior, *F*(2.428, 150.5) = 218.1, *p* < 0.0001, indicating that freezing increased across successive pairings, consistent with normal fear acquisition. In contrast, there was no significant main effect of treatment, *F*(2, 62) = 1.502, *p* = 0.23, and no treatment × CS-US presentation interaction *F*(4.856, 150.5) = 1.652, *p* = 0.15, confirming baseline equivalence across groups (Figure 1A).

**Figure 1:**
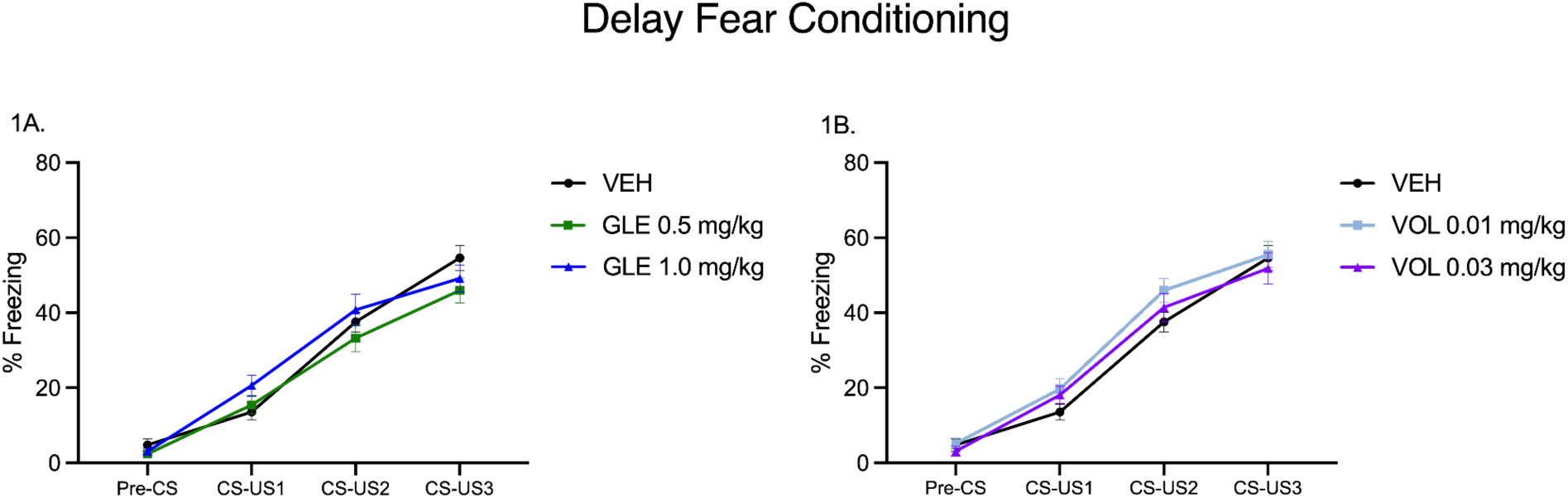
Baseline Fear Conditioning Across Future Treatment Groups. All mice underwent delay fear conditioning consisting of three tone-shock (CS-US) pairings. Freezing behavior increased across trials in all groups, confirming successful fear acquisition. (1A) Freezing behavior during delay fear conditioning in mice assigned to the MDL 11,939 cohort. A two-way RM ANOVA yielded a significant main effect of CS-US presentation (*p* < 0.0001), but no significant effect of treatment (*p* = 0.23) and no interaction (*p* = 0.15). (1B) Freezing behavior during delay fear conditioning in mice assigned to the MDL 100,907 cohort. A two-way RM ANOVA yielded a significant main effect of CS-US presentation (*p* < 0.0001), but no significant effect of treatment (*p* = 0.28) and no interaction (*p* = 0.53). Data are presented as mean ± SEM. Treatment groups: VEH (n=23), GLE 0.5 mg/kg (*n*=22), GLE 1.0 mg/kg (*n*=20), VOL 0.01 mg/kg (*n*=25), VOL 0.03 mg/kg (*n*=25).

Similarly, in the MDL 100,907 cohort (VEH, VOL 0.01 mg/kg, VOL 0.03 mg/kg), a two-way RM ANOVA revealed a significant main effect of CS-US presentation on freezing behavior, *F*(2.203, 154.2) = 244.1, *p* < 0.0001), with no main effect of treatment *F*(2, 70) = 1.303, *p* = 0.28 and no interaction *F*(4.405, 154.2) = 0.8107, *p* = 0.53, indicating equivalent baseline conditioning across groups (Figure 1B).

Together, these results indicate that all treatment groups acquired the conditioned freezing response prior to extinction training and testing

### 3.2. Freezing During First CS Presentation on Extinction Day 1

To assess whether MDL 11,939 or MDL 100,907 affected retrieval of conditioned fear memory, % freezing during the first CS presentation was analyzed using a one-way ANOVA within each cohort.

In the MDL 11,939 cohort, there was a significant main effect of treatment (*F*(2,62) = 3.71, *p* = 0.030). Dunnett’s post-hoc analysis revealed mice treated with 1.0 mg/kg exhibited significantly increased freezing compared to vehicle controls (*p =* 0.036), whereas the 0.5 mg/kg group did not differ from the vehicle group (*p =* 0.051) (Figure 2A). In the MDL 100,907 cohort, a significant main effect was also observed (*F*(2,70) = 3.18, *p =* 0.048). Dunnett’s comparison revealed increased freezing in the 0.01 mg/kg group compared to vehicle mice (*p =* 0.029), whereas the 0.03 mg/kg did not differ significantly (*p* = 0.164) (Figure 2B)

**Figure 2:**
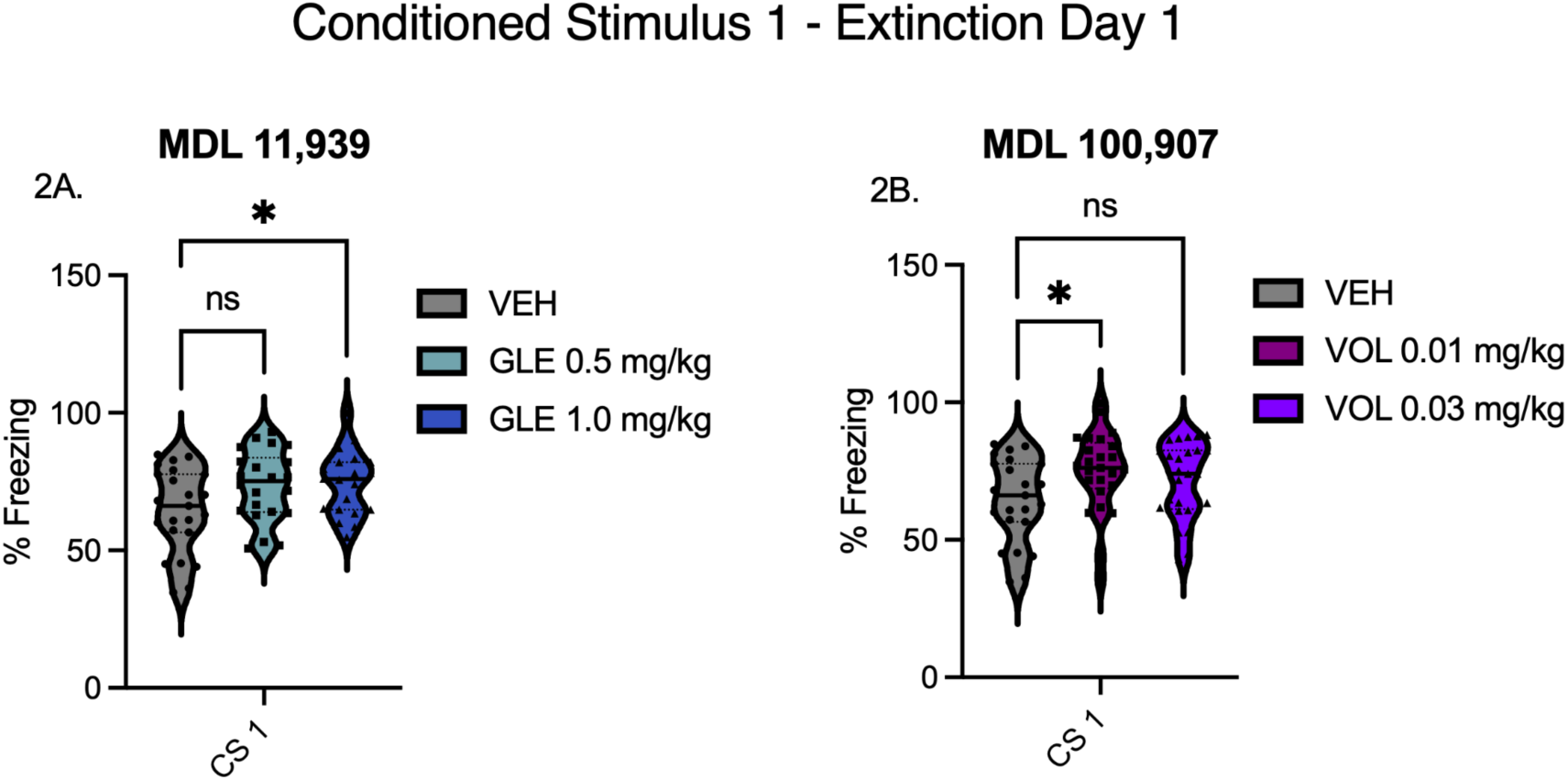
Freezing during the first CS presentation on Extinction Day 1. (2A) MDL 11,939 cohort. One-way ANOVA revealed a significant effect of treatment (*F(2,62) = 3.71, p* = 0.030), with the 1.0 mg/kg treated mice freezing more compared to VEH (Dunnett’s, *p =* 0.036) whereas the 0.5 mg/kg group did not differ from vehicles. (2B). MDL 100,907 cohort. One-way ANOVA revealed a significant effect of treatment (*F*(2,70) = 3.18, *p =* 0.048), with the 0.01 mg/kg treated mice having significantly increased freezing relative to vehicles (Dunnett’s, *p =* 0.029).

These findings suggest that acute 5-HT_2_AR antagonism can enhance the initial expression of conditioned fear, reflected by increased freezing during the first CS presentation on Extinction Day 1. MDL 11,939 producing a dose-related increase and MDL 100,907 exhibiting a significant effect but at the lower dosage.

### 3.3 Effects of MDL 11,939 on Acquisition and Retention of Fear Extinction

#### 3.3.1. Extinction Day 1 (Acute and Repeated Dose, First Dose)

A two-way RM ANOVA revealed a significant main effect of treatment, *F*(2, 62) = 3.34, *p* = 0.042, indicating overall group differences in freezing behavior. There was also a significant main effect of CS presentation, *F*(10.06, 623.5) = 37.67, *p* < 0.0001, reflecting a general decline in freezing across the session. However, the treatment × CS presentation interaction was not significant, *F*(20.11, 623.5) = 1.14, *p* = 0.30, suggesting that extinction rates were comparable across groups.

Tukey’s post-hoc tests did not yield significant pairwise differences between drug-treated groups and vehicle. A single comparison between GLE 0.5 mg/kg and GLE 1.0 mg/kg reached significance at CS13 (*p* = 0.024), suggesting a transient dose-related divergence in freezing behavior. However, this effect was not consistent across trials and did not differentiate drug groups from controls. Overall, these results suggest that a single dose of MDL 11,939 exerted a modest, dose-dependent influence on freezing behavior during the first extinction session, with higher doses generally associated with elevated freezing relative to vehicle (Figure 3A).

**Figure 3:**
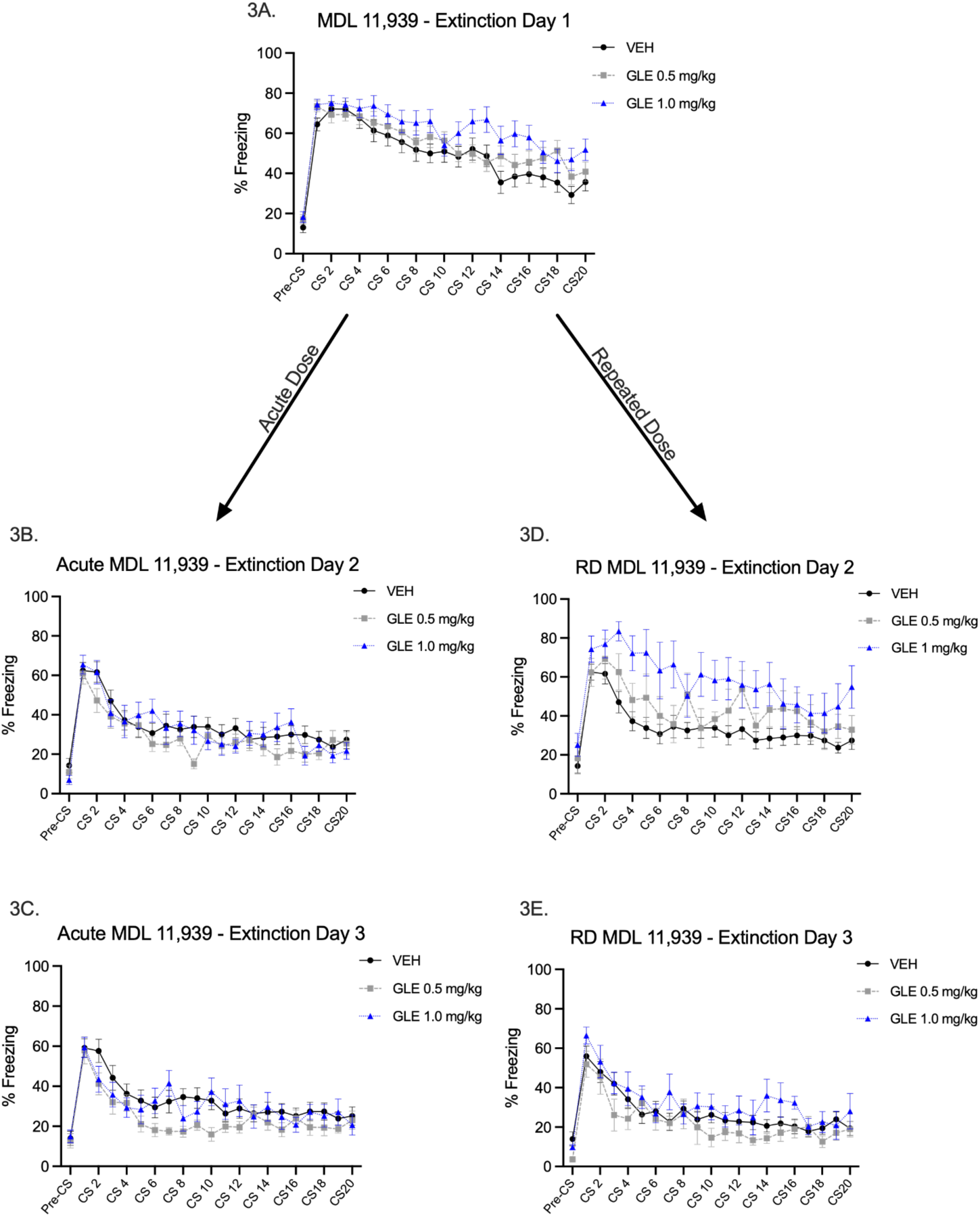
Acute and Repeated MDL 11,939 administration on fear extinction. (3A) Percent freezing across 20 conditioned stimulus presentations during Extinction Day 1 following acute administration of MDL 11,939 (0.5 mg/kg or 1.0 mg/kg). (3B-C) Twenty-four hours later mice were returned for Extinction Day 2 and 3, respectively to test retention of fear extinction. (3D) Twenty-four hours later mice were returned for Extinction Day 2 and administered a second dose of MDL 11,939 to test for freezing behavior following a repeated dose administration. (3E) Twenty-four hours after the repeated dose administration, mice were returned and tested drug-free to test retention of fear extinction.

#### 3.3.2 Extinction Day 2: Acute Dose

To assess the persistence of drug effects following a single dose of MDL 11,939, freezing behavior was analyzed during the second extinction session, conducted drug-free 24 hours after Extinction Day 1. Because a subset of mice transitioned into the repeated dosing protocol, sample sizes were reduced for the acute MDL 11,939 study: VEH (n=23), GLE 0.5 mg/kg (n=14) and GLE 1.0 mg/kg (n=12).

A two-way RM ANOVA revealed a significant main effect of CS presentation, *F*(20, 920) = 24.74, *p <* 0.0001, indicating continued extinction across trials. However, neither the main effect of treatment, *F*(2, 46) = 0.78, *p =* 0.46, nor the treatment × CS presentation interaction, *F*(40, 920) = 1.15, *p =* 0.29, reached significance. These results suggest that any drug-induced effects observed on Extinction Day 1 were not sustained 24 h later during the continuation of extinction learning (Figure 3B).

#### 3.3.3 Extinction Day 3: Acute Dose

To evaluate whether any residual effects of a single MDL 11,939 dose persisted into a second drug-free day, freezing behavior was analyzed during Extinction Day 3. Final group sizes were: VEH (*n*=23), GLE 0.5 mg/kg (*n* = 14) and GLE 1.0 mg/kg (*n* = 12). A two-way RM ANOVA revealed a significant main effect of CS presentation, *F*(10.38, 477.3) = 16.93, *p* < 0.0001, indicating continued extinction across trials. However, there was no significant main effect of treatment, *F*(2, 46) = 1.73, *p* = 0.19, and no treatment × CS presentation interaction, *F*(20.75, 477.3) = 1.19, *p* = 0.252. These findings suggest that the modest differences observed during earlier extinction sessions were no longer evident by Day 3 in the absence of the drug (Figure 3C).

#### 3.3.4 Extinction Day 2: Repeated Dose

To determine whether repeated administration of MDL 11,939 disrupts extinction learning, a subset of mice that received drug before Extinction Day 1 received the same treatment (VEH (*n*=23), GLE 0.5 mg/kg (*n*=8), or GLE 1.0 mg/kg (*n*=8) 30 min before Extinction Day 2. Freezing behavior was analyzed during each of 20 CS only presentations on Day 2.

A two-way RM ANOVA revealed a significant main effect of treatment, *F*(2, 36) = 5.37, *p* = 0.009, indicating a dose-dependent impairment of extinction. Mice treated with 1.0 mg/kg exhibited markedly elevated freezing across the session compared to vehicle controls. There was also a significant main effect of CS presentation, *F*(20, 720) = 13.36, *p* < 0.0001, reflecting extinction across all groups. However, the absence of a significant treatment × CS presentation interaction, *F*(40, 720) = 1.08, *p* = 0.36, suggests that the drug effect was consistent throughout the session, rather than trial-specific.

Tukey’s post-hoc analyses confirmed that mice in the 1.0 mg/kg MDL 11,939 froze significantly more than vehicle-treated controls beginning at CS3 (*p* = 0.0001), with additional group differences at CS4 (*p* = 0.014) and CS5 (*p* = 0.035). Although subsequent differences did not reach statistical significance, % freezing in the 1.0 mg/kg group remained numerically higher across the session than that of the vehicle and 0.5 mg/kg groups. These results suggest that repeated 5-HT_2_AR antagonism with MDL 11,939 impairs extinction learning, with the highest dose producing the most persistent disruption (Figure 3D).

#### 3.3.5 Extinction Day 3: Repeated Dose

To determine whether the effects of repeated MDL 11,939 administration persisted beyond the treatment window, freezing behavior was analyzed during Extinction Day 3, 24 h after the last injection. Final group sizes were: VEH (n=23), GLE 0.5 mg/kg (n=8) and GLE 1.0 mg/kg (n=8). A two-way repeated-measures ANOVA revealed a significant main effect of CS presentation, *F*(20, 720) = 15.07, *p <* 0.0001, indicating continued extinction across trials in all groups. However, there was no significant main effect of treatment, *F*(2, 36) = 1.48, *p* = 0.24, and no treatment × CS presentation interaction, *F*(40, 720) = 0.72, *p* = 0.79. These findings suggest that the elevated freezing observed with repeated dosing on Day 2 had dissipated by Day 3, and extinction proceeded at comparable rates across groups in the absence of the drug treatment (Figure 3E).

### 3.4 Effects of MDL 100,907 on Acquisition and Retention of Fear Extinction

#### 3.4.1. Extinction Day 1 (Acute and Repeated Dose, First Dose

Freezing behavior on Extinction Day 1 was analyzed to assess the acute effects of MDL 100,907. A two-way RM ANOVA revealed significant main effects of treatment group, *F*(2, 70) = 4.34, *p* = 0.017, and CS presentation, *F*(8.02, 561.2) = 26.51, *p* < 0.0001, confirming a general decline in freezing across trials (Figure 4A). The treatment × CS presentation interaction was not significant, *F*(38, 1330) = 0.66, *p* = 0.95, indicating that extinction rates were comparable across groups.

**Figure 4:**
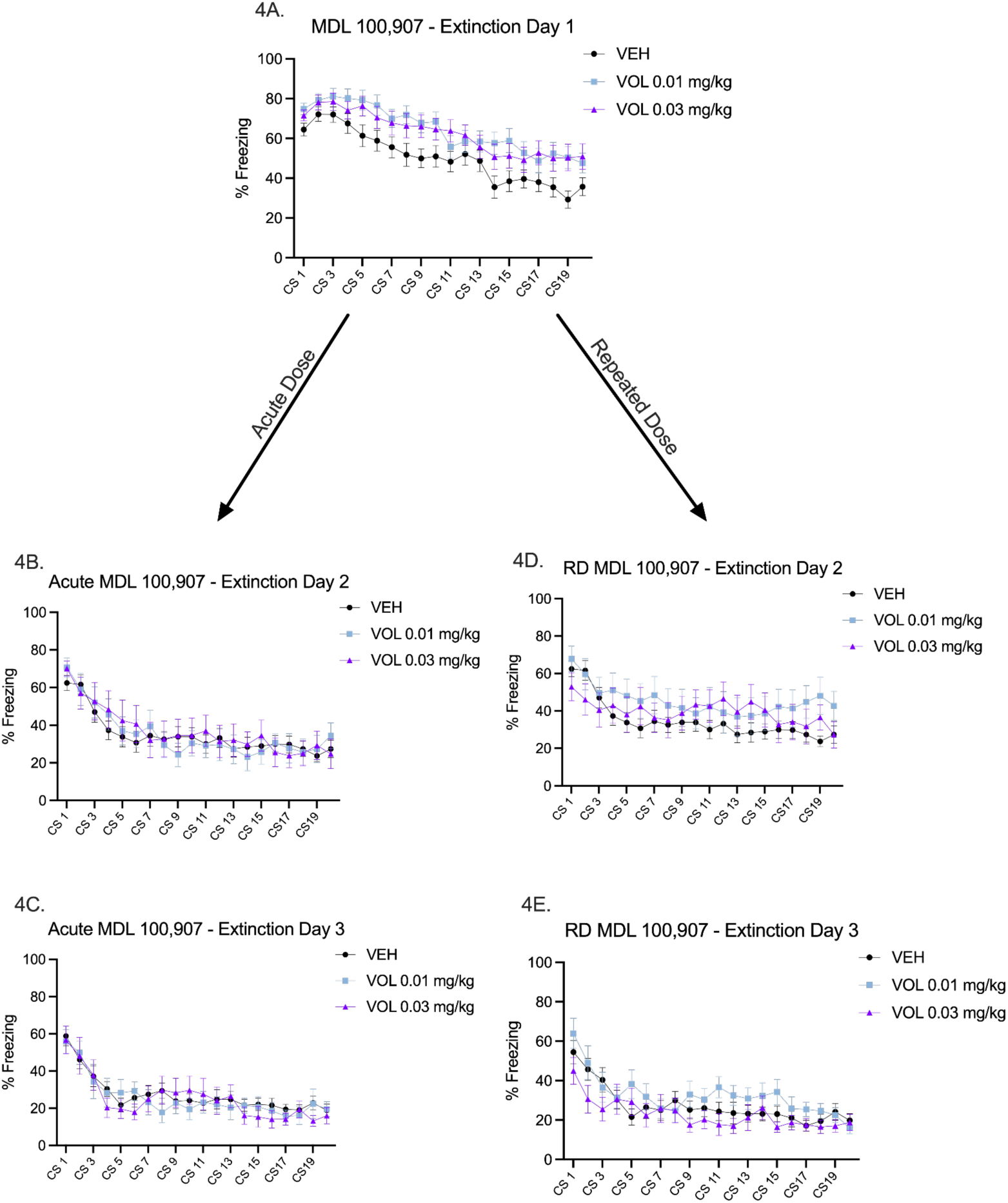
Acute and Repeated MDL 100,907 administration on fear extinction. (4A) Percent freezing across 20 conditioned stimulus presentations during Extinction Day 1 following acute administration of MDL 100,907 (0.01 mg/kg or 0.03 mg/kg). (4B-C) Twenty-four hours later mice were returned for Extinction Day 2 and 3, respectively to test retention of fear extinction (4D) Twenty-four hours later mice were returned for Extinction Day 2 and administered a second dose of MDL 100,907 to test for freezing behavior following a repeated dose administration. (4E) Twenty-four hours after the repeated dose administration, mice were returned and tested drug-free to test retention of fear extinction.

These effects were isolated to specific trials and were not maintained across the extinction session. Overall, MDL 100,907 produced subtle, transient increases in freezing during extinction, but did not alter the overall extinction trajectory. These findings suggest a modest impairment in extinction expression, rather than a robust or dose-dependent disruption of extinction learning.

#### 3.4.2 Extinction Day 2: Acute Dose

A two-way RM ANOVA revealed a significant main effect of CS presentation, *F*(8.38, 368.9) = 19.68, *p* < 0.0001, indicating a general reduction in % freezing across trials. Neither the main effect of treatment, *F*(2, 44) = 0.05, *p* = 0.95, nor the interaction between treatment and CS presentation, *F*(38, 836) = 0.69, *p* = 0.92, reached significance. These findings suggest that acute MDL 100,907 did not affect extinction behavior relative to vehicle controls (Figure 4B).

#### 3.4.3 Extinction Day 3: Acute Dose

A two-way RM ANOVA revealed a significant main effect of CS presentation, *F*(9.65, 424.8) = 19.65, *p* < 0.0001, indicating successful extinction across all groups. Neither the main effect of treatment, *F*(2, 44) = 0.11, *p* = 0.90, nor the treatment × CS presentation interaction, *F*(19.31, 424.8) = 0.76, *p* = 0.76, was significant. Thus, acute administration of MDL 100,907 did not affect % freezing levels on Day 3 of extinction (Figure 4C).

#### 3.4.4 Extinction Day 2: Repeated Dose

A two-way RM ANOVA on freezing measures from Extinction Day 2 of the repeated dosing study revealed a significant interaction between treatment and CS presentation, *F*(38, 874) = 1.56, *p* = 0.0177, and a significant main effect of CS presentation, *F*(10.51, 483.5) = 8.83, *p* < 0.0001, consistent with extinction across groups. The main effect of treatment was not significant, *F*(2, 46) = 0.86, *p* = 0.43. Despite the interaction, post-hoc Tukey’s tests did not reveal any significant pairwise differences between treatment groups. These results suggest subtle, session-dependent variation in freezing across groups, but no robust treatment effect (Figure 4D).

#### 3.4.5 Extinction Day 3: Repeated Dose

A two-way RM ANOVA on freezing measures from Extinction Day 3 of the repeated dosing study revealed no significant main effect of treatment, *F*(2, 42) = 1.39, *p* = 0.26, and no significant treatment × CS presentation interaction, *F*(38, 798) = 1.18, p = 0.21. The main effect of CS presentation was significant, *F*(9.55, 401.1) = 13.40, *p* < 0.0001, confirming extinction across groups. Thus, repeated administration of MDL 100,907 did not significantly alter % freezing behavior on Extinction Day 3 (Figure 4E).

### 3.5 Trials-to-Extinction

To evaluate overall differences in the acquisition of fear extinction, the number of trials required to reach extinction criterion were analyzed across treatment conditions using a one-way ANOVA. A significant main effect of treatment was observed, *F*(8, 106) = 3.36, *p* = 0.0018, indicating that drug treatment significantly influenced extinction learning. Tests for homogeneity of variance suggested unequal group variances (Brown-Forsythe: *F*(8, 106) = 3.46, *p* = 0.0014; Bartlett’s: *p* = 0.0095). Planned comparisons were restricted to treatment groups versus vehicle controls, Dunnett’s multiple comparisons test was retained for post hoc analysis (see Figure 5).

**Figure 5.**
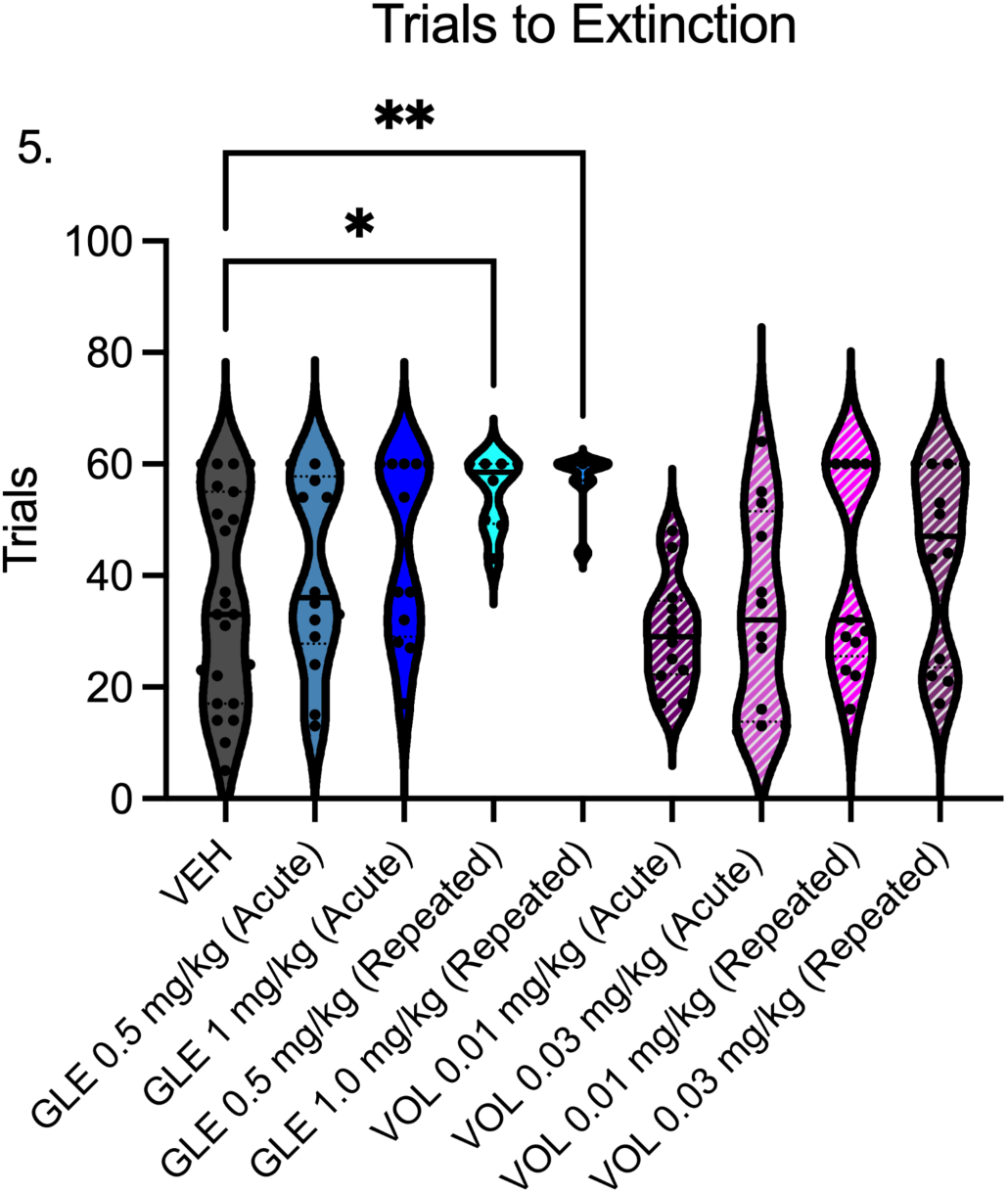
Trials to reach extinction criterion. Number of trials required to reach the extinction criterion defined as 4 or more consecutive CS presentations in which the % freezing response reached ≤50% of that during the third CS-US pairing on conditioning day. A one-way ANOVA revealed a significant effect of treatment F(8, 106) = 3.37, p = 0.0018). Dunnett’s post hoc comparisons comparing MDL 11,939- or MDL 100,907-treated mice to vehicles indicated that repeated administration of MDL 11,939 at 0.5 mg/kg (p = 0.026) and 1.0 mg/kg (p = 0.008) significantly increased the number of trials required to reach extinction. No other treatment groups differed from vehicles. Data are presented as mean ± SEM. Increased trial numbers reflect impaired extinction learning.

Dunnett’s multiple comparisons test (relative to vehicle) revealed that repeated administration of MDL 11,939 significantly increased the number of trials needed to reach extinction criterion at both 0.5 mg/kg (p = 0.026) and 1.0 mg/kg (p = 0.008), consistent with impaired extinction learning. In contrast, none of the MDL 100,907 treatment groups differed significantly from vehicle (all p > 0.05).

Together, these findings indicate that repeated dosing with MDL 11,939, a non-selective 5-HT2A/2C receptor antagonist, impaired extinction learning, as evidenced by increased freezing and resistance to extinction. By comparison, the selective 5-HT2A antagonist MDL 100,907 did not significantly alter the acquisition or retention of fear extinction (Figure 5).

## 4. Discussion

Antipsychotic medications are known to target the brain receptors that induce hallucinations (Peng et al., 2024). Most currently approved antipsychotic medications either partially or completely inhibit D2- and 5-HT_2_AR. Both receptors are essential for fear extinction, and fear is strongly associated with psychotic experiences. The therapeutic potential of antipsychotics would be limited if they alleviate hallucinations and other psychotic symptoms while aggravating fear-related psychopathy. Clinical studies of the selective 5-HT_2_A antagonists, MDL 11,939 and MDL 100,907 yielded clinically insignificant antipsychotic efficacy. MDL 11,939 was also clinically tested on patients with generalized anxiety disorder, but failed to improve symptoms (Sramek et al., 1995), and while MDL 100,907 has not been tested clinically, it has shown no significant impact on depression-like behavior or insomnia in rodents (Marek et al., 2005; Teegarden et al., 2008).

Despite their lack of clinical efficacy, selective antagonists offer an opportunity to examine fear extinction under conditions of 5-HT_2_A inhibition, a pharmacological property shared by many currently prescribed antipsychotics. In this study, we investigated the effects of systemic administration of MDL 11,939 and MDL 100,907 on the acquisition and retention of fear extinction in adult C57BL/6J mice. MDL 11,939 produced modest but dose-dependent impairments in extinction, particularly after repeated dosing, as evidenced by persistently elevated freezing during extinction sessions and a significantly greater number of trials required to meet extinction criteria, relative to vehicle-treated controls. Fear extinction remained impaired with a repeat exposure to MDL 11,939, consistent with prior reports showing that this compound does not induce 5-HT_2_A overexpression in mice. If repeated MDL 11,939 administration had upregulated the 5-HT_2_A receptor, enhanced serotonergic signaling may have been expected to facilitate extinction learning. The persistence of extinction deficits suggests that impaired extinction may not be driven by compensatory receptor upregulation, but rather by sustained antagonism or ligand-specific signaling at the 5-HT_2_A receptors. Although MDL 100,907 increased freezing during the first CS presentation on Extinction Day 1, indicating enhanced expression of conditioned fear, it did not produce persistent impairments in extinction learning or retention across subsequent extinction sessions. Because the drugs were administered systemically, the effects observed with MDL 11,939 likely reflect receptor antagonism across multiple brain regions involved in extinction, including the prelimbic cortex, infralimbic cortex, and amygdala, rather than action within a single region. Taken together, the absence of an effect with the more selective antagonist MDL 100,907 suggests that selective 5HT_2_A receptor blockade does not meaningfully disrupt fear extinction under the conditions tested.

5-HT_2_A receptors can signal in very different ways depending on how they are activated, and these differences matter for behaviors (psychedelic effects vs. learning vs. fear extinction). Biased 5-HT_2_A ligands differ in their selectivity to each of the receptor intracellular signaling pathways, which include G_q/11_ activating PLC, β-arrestin desensitizing the receptor to G-proteins and activating ERK/MAPK pathway, G_i_ inhibiting adenylyl cyclase, G_12/13_ activating PLA2, and ARF activating phospholipase D (Jastrzębski et al., 2025). In mice, agonists that stimulate the G_i1_ pathway induced a head twitch behavior - a manifestation of psychedelic effects - whereas agonists of the G_q_ pathway suppressed formation of long-term memory (Kossatz et al., 2024). Non-hallucinogenic agonists are biased to the G_q_-dependent signaling, and psychedelics frequently activate G_i1_- and β-arrestin-2-dependent pathways. Because a combination of an SSRI and 5-HT_2_A inverse agonist enhances fear extinction without head twitch behavior, G_i1_ pathway may not be essential for fear extinction.

When MDL 100,907 and MDL 11,939 were tested in mice and post-mortem human brains for their ability to modify G_i1_ and G_q/11_ pathways, MDL 100,907 suppressed both pathways, whereas MDL 11,939 showed no effect and blocked the effects of other tested ligands (Muneta-Arrate et al., 2025). Therefore, MDL 11,939 is an antagonist and MDL 100,907 is inverse agonist in respect to the G_i1_ and G_q/11_ signaling pathways. This implies that 5-HT_2_A receptors are constitutively active, so inverse agonists block stimulation, and silence ongoing signaling. Furthermore, MDL 11,939 and MDL 100,907 differ in their ability to weaken facilitation of fear extinction usually produced by psychedelic compounds. DOI is a classical serotonergic psychedelic with affinity to multiple 5-HT2 receptor subtypes, whereas TCB-2 is a more selective 5-HT2AR agonist (Poulie et al., 2022). In contrast, 25CN-NBOH is a highly selective 5-HT2AR agonist acting on β-arrestin-2-biased signaling. Compared to DOI, its reported head-twitch behavior in mice was consistent with partial agonism, and 25CN-NBOH inhibited the DOI effect in dose-dependent manner when combined with DOI (Fantegrossi et al., 2015). In contrast, 25CN-NBOH was identified as the first β-arrestin-2-biased 5-HT_2_A agonist (Poulie et al., 2022). Compared to DOI, its reported effect on head twitch behavior in mice was consistent with partial agonism, and 25CN-NBOH inhibited the DOI effect in dose-dependent manner when combined with DOI (Fantegrossi et al., 2015). MDL 11,939 blocked the memory-enhancing effect of TCB-2 (Zhang et al., 2013), whereas MDL 100,907 blocked the reduction in conditioned fear response induced by 25CN-NBOH, but had no impact on the freezing-reducing effect of TCB-2 (Hagsäter et al., 2021). Therefore, MDL 11,939 and MDL 100,907 may have different molecular mechanisms by which 5-HT_2_A antagonism is achieved. Although the effects of MDL 11,939 on β-arrestin pathway are not known, antagonism of serotonin stimulation of the β-arrestin pathway may contribute to impaired fear extinction. Serotonin can activate the β-arrestin pathway and was shown to induce head twitch behavior in mice by a β-arrestin-2-dependent mechanism (Schmid et al., 2008). The difference in behavioral effects between MDL 11,939 and MDL 100,907 suggest that fear extinction facilitation may depend on an existing molecular mechanism that is susceptible to inhibition by MDL 11,939 but not MDL 100,907. TCB-2 rather than 25CN-NBOH may activate this mechanism through 5-HT_2_A because the selective 5-HT_2_A antagonist MDL 11,939 blocks the TCB-2 effect (Zhang et al., 2013). β-arrestin may not mediate this presumptive mechanism because β-arrestin-biased agonist 25CN-NBOH generates about half the TCB-2 effect, and MDL 100,907 can block the 25CN-NBOH effect but not the TCB-2 effect (Hagsäter et al., 2021; Poulie et al., 2022). We hypothesize two distinct mechanisms that may facilitate fear extinction: one dependent on the β-arrestin pathway and another susceptible to inhibition by MDL 11,939. Boosting the extracellular serotonin concentration with an SSRI and inhibition of competing intracellular pathways with MDL 100,907 or pimavanserin is more consistent with activation of the latter mechanism, which remains elusive but may involve stimulation of intracellular 5-HT_2_A receptors. This interpretation is supported by evidence that nonhallucinogenic 5-HT_2_A agonists stimulate predominantly the G_q/11_ pathway without any impact on fear extinction. Together these findings suggest that modulating the G_q/11_ pathway signaling alone may not be sufficient to explain the behavioral differences observed between these two receptor ligands in the present study.

Consistent with the complexity of 5-HT_2_A receptor signaling, an *in-vitro* study compared the effective doses of multiple established antipsychotics with their affinity to cloned D2, 5-HT1A, 5-HT_2_A, 5-HT_2_C receptors, reporting a strong correlation between D2 receptor affinity and antipsychotic potency, whereas the 5-HT_2_C and 5HT_1_A binding affinities demonstrated weaker and mostly inverse correlation (Richtand et al., 2007). Notably, there was no correlation between antipsychotic potency and the 5-HT_2_A affinity, potentially reflecting the diversity of intracellular signaling pathways of the 5-HT_2_A receptor, rather than a lack of functional relevance. The existence of multiple biased profiles of 5-HT_2_A receptor antagonists and inverse agonists indicate a possibility that impaired fear extinction may represent an underrecognized adverse effect of antipsychotics. These effects would not be expected to scale with potency but instead depend on ligand-specificity and dosing regimen.

The exclusive use of systemic drug administration enhances translational relevance but limits anatomical specificity. Consequently, the impairments observed with MDL 11,939, particularly with repeated dosing, are likely due to network-level consequences of broad 5-HT_2_ receptor antagonism. Because MDL 11,939 antagonizes both 5-HT_2_A and 5-HT_2_C receptors, its effects likely engage multiple receptor populations across distinct brain regions, complicating attribution to a single mechanism or site of action. To evaluate whether 5-HT_2_C receptor antagonism contributed to the observed extinction impairments seen by repeated MDL 11,939 administration, we assessed the effects of selective 5-HT_2_C manipulation in a cohort of naive mice, and observed no significant impact on fear acquisition, extinction or trials-to-extinction (Supplementary Fig. S1). Therefore, selective antagonism of the 5-HT_2_C receptor does not recapitulate the deficits in extinction learning that is observed with repeated MDL 11,939 administration, suggesting that 5-HT_2_C receptor blockade alone is not sufficient to replicate these effects.

Future studies combining systemic and localized intracranial pharmacological manipulations, receptor occupancy mapping, and molecular or circuit-level readouts (e.g., immediate early gene expression, electrophysiology, or calcium imaging) will be essential to identifying the receptor and circuit mechanisms underlying our observed behavioral effects. Despite these limitations, the present findings demonstrate that fear extinction deficits may represent an underacknowledged behavioral consequence of serotonergic antagonism. Such an effect may not be predicted by antipsychotic potency or receptor affinity alone but instead emphasizes ligand-specific signaling and dosing history that considers intracellular signaling mechanisms when evaluating cognitive and behavioral effects of serotonergic therapies.

## Supplementary Results

### S1. Assessment of 5-HT_2_C Receptor Involvement in Fear Extinction

To determine whether the 5-HT_2_C receptor contributes to the effects of MDL 11,939 on fear extinction behavior in the main study, a naive cohort of adult C57BL/6J mice underwent auditory fear conditioning, drug-free, followed by extinction training across three consecutive days after systemic administration of vehicle or the selective 5-HT_2_C receptor antagonist, SB 242084 (0.25 or 0.50 mg/kg).

During conditioning, freezing behavior increased across CS–US pairings, reflected by a significant main effect of CS presentation (two-way RM ANOVA, *p* < 0.0001), with no main effect of treatment (*p* = 0.2884) and no CS × treatment interaction (*p* = 0.5683; Supplementary Fig. S1A).

To test whether selective 5-HT_2_C receptor antagonism influenced retrieval of conditioned fear, freezing during the first CS presentation on Extinction Day 1 was analyzed using a one-way ANOVA. No significant effect of treatment was observed *F*(2,25) = 0.02, *p =* 0.981. Dunnett’s post hoc comparisons relative to vehicles revealed no differences between SB 242084 treated mice (0.25 mg/kg or 0.5 mg/kg) and vehicle controls (all *p >* 0.98) (Supplementary Fig. S1B). These findings indicate that selective 5-HT_2_C receptor antagonism does not alter initial freezing during the first extinction test.

During extinction training, freezing responses declined across CS presentations on Extinction Days 1–3, as indicated by significant main effects of CS presentation on each day (*p* < 0.0001). However, no main effects of treatment (Day 1: *p* = 0.6158; Day 2: *p* = 0.9786; Day 3: *p* = 0.3657) and no CS × treatment interactions (all *p* > 0.44) were observed (Supplementary Fig. S1C–E).

Consistent with these findings, the number of trials required to reach extinction criteria did not differ between treatment groups (one-way ANOVA, *p* > 0.05; Tukey’s multiple comparisons, all adjusted *p* > 0.30; Supplementary Fig. S1F).

Together, these data indicate that selective 5-HT_2_C receptor blockade did not significantly alter fear acquisition or extinction learning under the conditions tested and is therefore unlikely to account for the extinction impairments observed following MDL 11,939 administration.

---

**Supplementary Figure S1.**
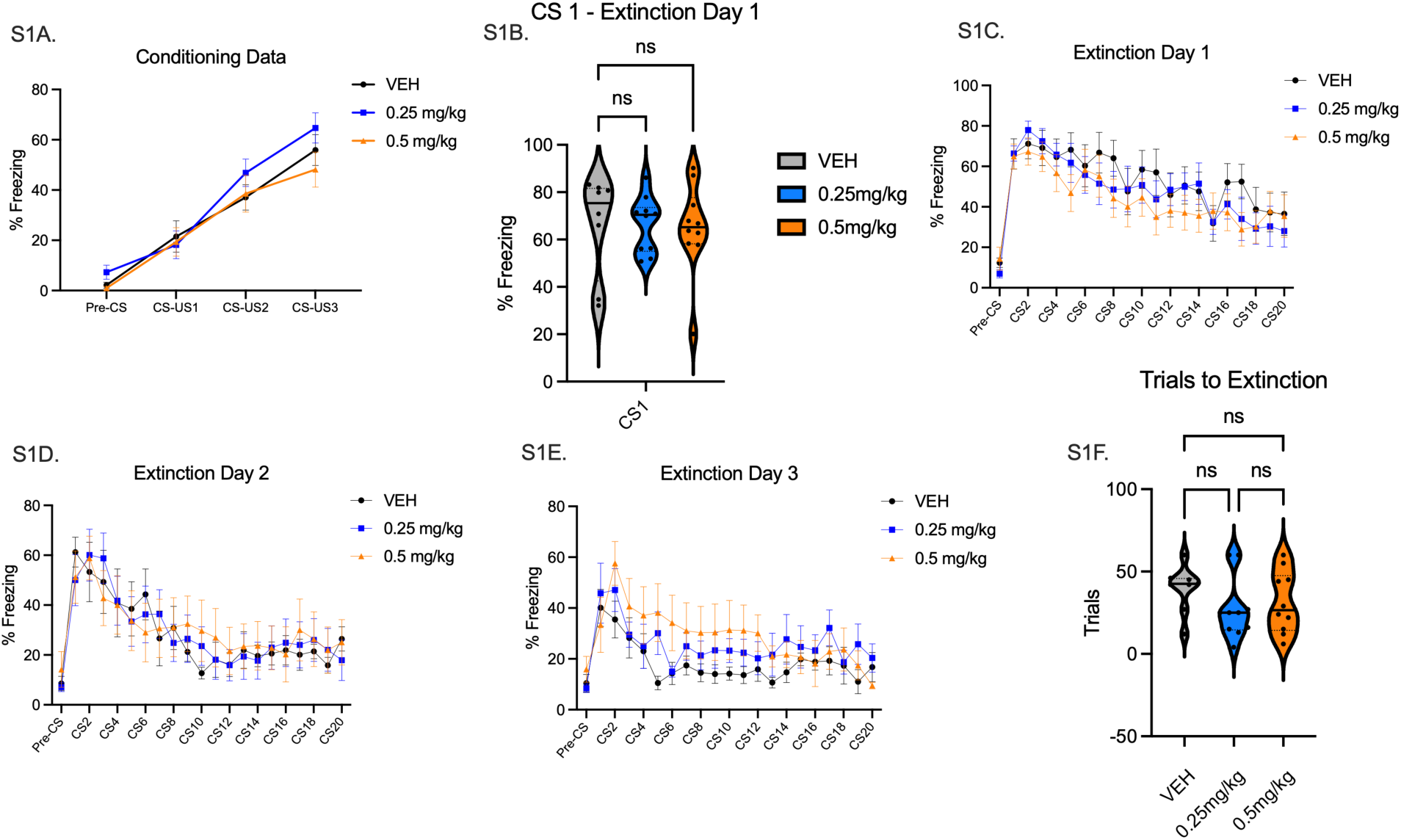
Selective 5-HT_2_C receptor blockade does not alter fear expression or the acquisition of fear extinction. (A) Mean percent freezing across future treatment groups during auditory fear conditioning with three CS–US pairings. (B) Freezing during the first CS presentation on Extinction Day 1. (C–E) Percent freezing in response to CS presentations during training on Extinction Days 1–3. (F) Number of trials required to reach the extinction criteria. Data are presented as mean ± SEM. Conditioning and extinction data were analyzed using two-way RM ANOVA with CS presentation as the within-subjects factor and treatment as the between-subjects factor, followed by Geisser–Greenhouse correction where appropriate. Trials-to-extinction were analyzed using one-way ANOVA with Tukey’s multiple comparisons test. No significant effects of treatment or CS × treatment interactions were detected.

